# Glucose promotes resistance of human commensal *Escherichia coli* against contact-killing by pandemic *Vibrio cholerae*

**DOI:** 10.1101/2020.06.22.166306

**Authors:** Cristian V Crisan, Holly Nichols, Sophia Wiesenfeld, Gabi Steinbach, Peter J Yunker, Brian K Hammer

## Abstract

Evolutionary arms races among organisms are broadly prevalent and bacteria have evolved defensive strategies against various attackers. A common microbial aggression mechanism is the Type VI Secretion System (T6SS), a contact-dependent bacterial weapon used to deliver toxic effector proteins into adjacent target cells. Sibling cells constitutively express immunity proteins that neutralize effectors. However, less is known about mechanisms that allow non-sibling bacteria to respond to external cues and survive T6SS attacks independently of immunity proteins. In this study, we show that resistance to T6SS attacks is promoted by a genetically controlled response to exogenous glucose. We observe that multiple human *Escherichia coli* commensal strains lacking immunity proteins are sensitive to T6SS attacks from pandemic *Vibrio cholerae* on nutrient-rich media. By contrast, *E. coli* cells become resistant to attacks when co-cultured on the same media with glucose. We confirm that glucose does not impair *V. cholerae* T6SS activity. Instead, we find that cAMP receptor protein (CRP), which alters expression of hundreds of genes in response to glucose, controls resistance to T6SS attacks in *E. coli* cells. Consistent with the observed resistance on media with glucose, an *E. coli crp* disruption mutant survives significantly better against *V. cholerae* T6SS attacks even in the absence of glucose. Finally, we also show that resistance to T6SS attacks depends on the pH of the medium and varies based on the target and killer strains.

**IMPORTANCE:** Many Gram-negative bacteria, including important pathogens, encode T6SS genes to deliver toxic effectors and eliminate competitors. Our results uncover a novel defense mechanism against T6SS attacks that is triggered by an external stimulus and mediated by a metabolic response in non-kin target cells. In microbiomes such as those in gastrointestinal tracts where T6SS activity is known to occur, signaling by metabolites like glucose may affect the efficacy of T6SS attacks and alter microbial community composition. Our findings could have vast implications for microbial interactions during pathogen colonization of hosts and survival of bacterial cells in environmental communities. Furthermore, the glucose-mediated resistance observed here might provide a novel example of an evolutionary arms race between killer T6SS cells and target bacteria.

## INTRODUCTION

*Vibrio cholerae* is the waterborne enteric pathogen that causes serious, often fatal cholera diarrheal disease when ingested by humans. This ubiquitous microbe is found in dense polymicrobial marine communities on chitinous surfaces and in animal reservoirs like fish or zooplankton (1–3). To compete with other cells in densely-populated microbial environments, *V. cholerae* employs a harpoon-like structure called the Type VI Secretion System (T6SS) (4–7). The T6SS punctures adjacent cells and delivers toxic effector proteins that disrupt lipid membranes and cell walls (8–10). In animal models, *V. cholerae* uses T6SS effectors to eliminate commensal bacteria like *Escherichia coli* and *Aeromonas veronii* (11, 12). *V. cholerae* strains with an active T6SS exhibit increased pathogenicity and cause severe cholera-like symptoms in animal model systems (11–13). These observations suggest the apparatus plays important roles in enhancing the ability of *V. cholerae* to colonize environmental habitats and infect hosts. However, while many studies have investigated T6SS offensive abilities, less is known about mechanisms of protection against T6SS aggression.

Bacterial cells with active T6SSs constitutively express immunity proteins that recognize and neutralize toxic effectors (9, 10, 14). The survival of *V. cholerae* cells devoid of immunity proteins can be modestly increased (~5-fold) by upregulating expression of secreted exopolysaccharides, presumably by creating a physical barrier between the killer and target strains (15). *Pseudomonas aeruginosa* does not possess cognate immunity proteins for *V. cholerae* T6SS effectors but is able to survive attacks through an unknown mechanism (16). Proteases, reactive oxygen species and membrane stress response systems modulate survival of target cells against T6SS attacks (17–20).

Previous studies found that environmental conditions and small molecules can affect the outcome of T6SS-mediated antagonism (20–22). Also, glucose has been shown to decrease the ability of human commensal bacteria to colonize intestinal tracts and alter interactions between *V. cholerae* and *E. coli* cells in hosts (23–25). Here we report that exogenous glucose allows human commensal *E. coli* strains to resist T6SS-mediated aggression from pandemic killer *V. cholerae* cells. We determine the resistance is dependent on the target and killer strains, the pH of the medium and the *V. cholerae* T6SS effectors. Finally, we demonstrate that protection against T6SS attacks is mediated by the CRP glucose response regulator.

## RESULTS

### Glucose allows *E. coli* cells to survive T6SS attacks

When co-cultured on rich LB medium, *V. cholerae* C6706 cells expressing the QstR protein constitutively (denoted here as C6706*) kill *E. coli* MG1655 target cells efficiently (Fig. 1A) (26–29). By contrast, a T6SS-*V. cholerae* C6706* strain with a deletion in the essential T6SS protein VasK is unable to eliminate target cells (Fig. 1A) (30). We observed that when *V. cholerae* C6706* and *E. coli* MG1655 cells are co-cultured on LB medium containing 0.4% glucose (LBG), survival of *E. coli* cells significantly increased (~1000 fold) compared to co-cultures on LB with no added glucose (Fig. 1A). The number of recovered *V. cholerae* killer cells was unaltered when co-cultured for three hours with *E. coli* MG1655 on LB and LBG (Fig. 1B).

**Figure 1.**
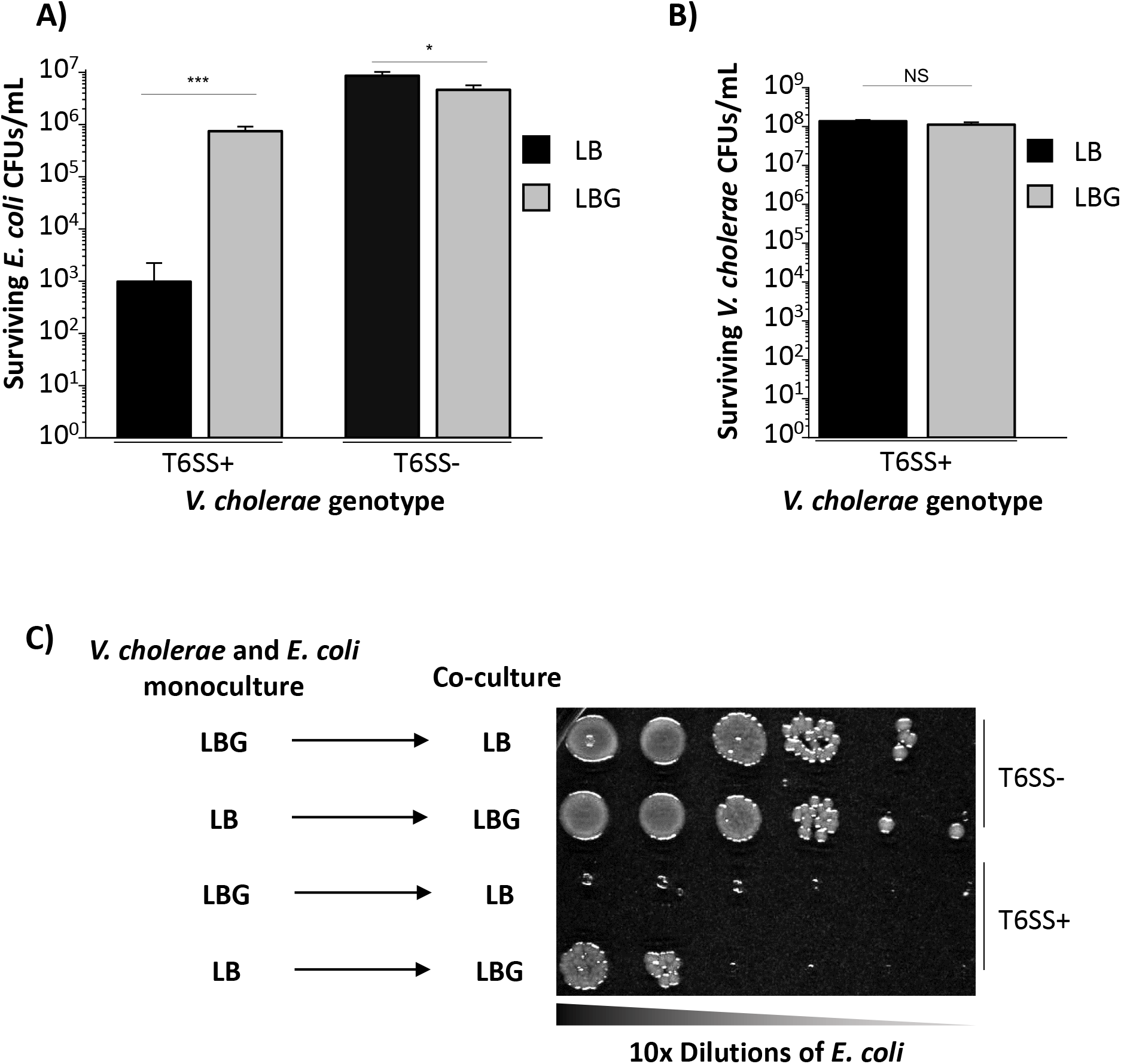
Glucose confers *E. coli* cells resistance against *V. cholerae* T6SS attacks. A) *E. coli* MG1655 cells were co-cultured for 3 hours with either killer T6SS+ or defective T6SS-*V. cholerae* C6706* cells on LB or LBG plates. The number of surviving *E. coli* cells was determined by counts of colony forming units (CFUs). A Student’s t-test was performed to determine significance. *** p < 0.001, * p < 0.05. B) The same competition assay was performed as described above. However, the number of surviving killer T6SS+ *V. cholerae* C6706* cells was determined instead. A Student’s t-test was performed to determine significance. NS not significant. C) *V. cholerae* C6706* cells were grown overnight in LB or LBG and then co-cultured with *E. coli* on LBG or LB, respectively.

To determine whether *E. coli* cells require active glucose induction to survive T6SS attacks, we incubated *V. cholerae* C6706* and *E. coli* MG1655 in liquid LBG or LB and then co-cultured the strains on either LB or LBG, respectively (Fig. 1C). *E. coli* cells resisted T6SS attacks when incubated in liquid LB and co-cultured with *V. cholerae* C6706* on LBG but remained susceptible when incubated in liquid LBG and co-cultured on LB (Fig. 1C). We also hypothesized that glucose could allow *E. coli* cells to escape T6SS attacks by replicating faster and evading killer cells. When grown individually in monoculture conditions that mimicked co-culture assays, growth rates of *E. coli* and *V. cholerae* were unaltered on LBG compared to LB (Supp. Fig. 1).

### Survival of target cells is dependent on the target strain, the sugar type and the killer strain

To test whether the robust glucose-mediated protection was specific to *E. coli* MG1655, we co-cultured killer *V. cholerae* C6706* with other human *E. coli* commensal strains (Nissle, HS and ECOR-2), as well as fish symbiotic *A. veronii* and susceptible *V. cholerae* (12, 31–37). We observed that all tested target strains had increased survival on LBG compared to LB medium (Fig. 2A). On LBG, target *V. cholerae* and *A. veronii* cells were still killed in relatively high numbers compared to *E. coli*, as the survival difference on LBG compared to LB was considerably larger for *E. coli* strains (greater than one order of magnitude) than for *V. cholerae* or *A. veronii* (less than or equal to one order of magnitude) (Fig. 2A).

**Figure 2.**
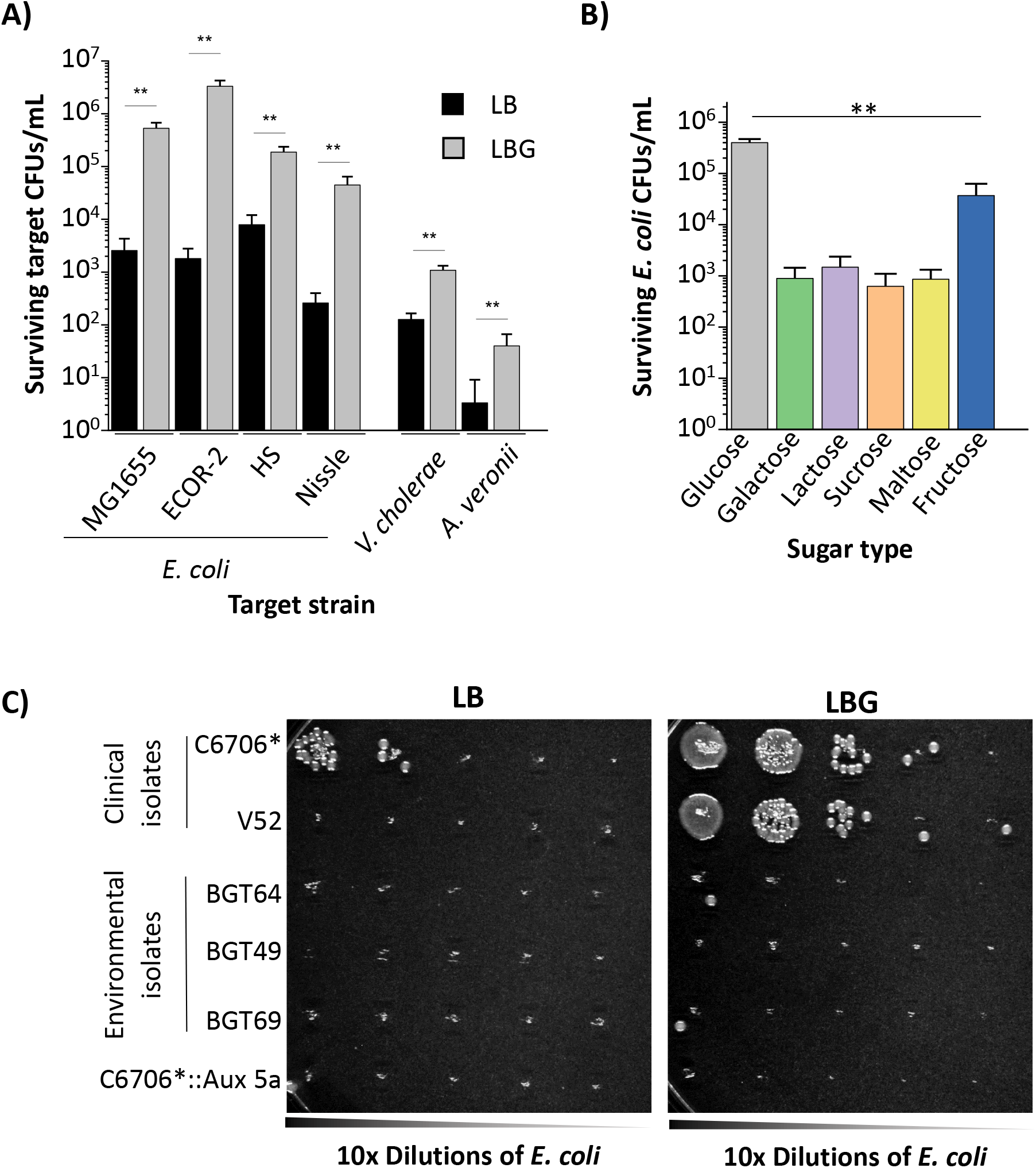
Resistance against T6SS attacks depends on the target strain, the killer strain and the sugar substrate. A) Along with MG1655, *E. coli* commensal strains Nissle, HS, ECOR-2, *A. veronii* and susceptible *V. cholerae* target cells were co-cultured with killer T6SS+ *V. cholerae* C6706* on LB or LBG. Student’s t-tests were performed to determine significance. ** p < 0.01, * p < 0.05. B) Killer *V. cholerae* C6706* cells were co-cultured with *E. coli* MG1655 cells on LB medium containing 0.4% of the indicated sugar compounds. A One-Way ANOVA with a post-hoc Tukey HSD test was used to determine significance. ** p < 0.01. C) Clinical killers *V. cholerae* C6706*, C6706*::Aux 5a and V52 as well as environmental *V. cholerae* strains BGT49, BGT64 and BGT69 strains were co-cultured with *E. coli* MG1655 cells on LB medium (left) or LBG medium (right).

We next investigated if the *E. coli* resistance to T6SS attacks is specific to glucose. We co-cultured killer C6706* and target *E. coli* MG1655 on LB medium containing different sugars: fructose and galactose (monosaccharides), as well as sucrose, maltose and lactose (glucose-containing disaccharides). The concentration for each sugar was 0.4%, as with glucose. All sugars resulted in significantly less survival of target cells when compared to glucose, with fructose conferring intermediate levels of protection (see Discussion). When identical co-culture experiments were conducted using T6SS-defective killer C6706* cells, the number of recovered *E. coli* MG1655 cells was similar for all tested sugars (Supp. Fig. 2).

We also sought to determine whether the effects observed were restricted to T6SS attacks by some strains of *V. cholerae* but not others. *V. cholerae* strain V52 is a clinical isolate that constitutively expresses T6SS genes on LB rich medium, yet encodes the same toxins as C6706 (10, 38, 39). Strains BGT49, BGT64 and BGT69 are *V. cholerae* isolates obtained from sources other than patients (referred to as environmental isolates) and display robust contact killing on LB medium (Fig. 2C) (40). These strains encode a diverse repertoire of T6SS toxins (41). On LBG medium, both V52 and C6706* killed fewer *E. coli* MG1655 cells than on LB (Fig. 2C). By contrast, strains BGT49, BGT64 and BGT69 bypassed the glucose-mediated protection and eliminated *E. coli* cells with similar efficacy on both LB and LBG (Fig. 2C). We reported that strain BGT49 encodes an additional toxin (*tleV1*) within the Aux 5a T6SS gene cluster that is absent from patient-derived strains like C6706 or V52 (41). We previously integrated the Aux 5a gene cluster onto the chromosome of *V. cholerae* C6706* T6SS+ (C6706*::Aux 5a) and showed that C6706*::Aux 5a can kill parental *V. cholerae* C6706 cells lacking the cognate immunity protein for TleV1 (41). When co-cultured with *E. coli* cells on LBG, *V. cholerae* C6706*:: Aux 5a bypassed the glucose-mediated protection and successfully eliminated *E. coli* cells, by contrast with C6706* (Fig. 2C).

### Buffered LBG media at a near-neutral pH significantly increases T6SS killing

When grown aerobically on glucose, *E. coli* MG1655 secretes organic acids that inhibit growth of *V. cholerae* cells (24, 25). Since we did not observe a decrease in the number of *V. cholerae* cells when co-cultured with *E. coli* cells on LBG after 3 hours, we hypothesized that the killing efficiency of *V. cholerae* could be affected by a pH change (Fig. 1B, Fig. 3A). When co-cultures of *E. coli* MG1655 and *V. cholerae* C6706* were performed on LBG medium buffered to a pH of 7.4, the number of surviving *E. coli* MG1655 cells was significantly decreased compared to unbuffered LBG medium (Fig. 3A). To determine if *E. coli* reduces the efficiency of T6SS-mediated killing on LBG by impairing delivery of *V. cholerae* effectors, we performed a polyculture assay in which *V. cholerae* C6706* T6SS+ killer cells were cultured with both *E. coli* and susceptible *V. cholerae* at the same time (Fig. 3B). While *E. coli* cells resisted T6SS attacks as expected, target *V. cholerae* cells were efficiently killed (Fig. 3B).

**Figure 3.**
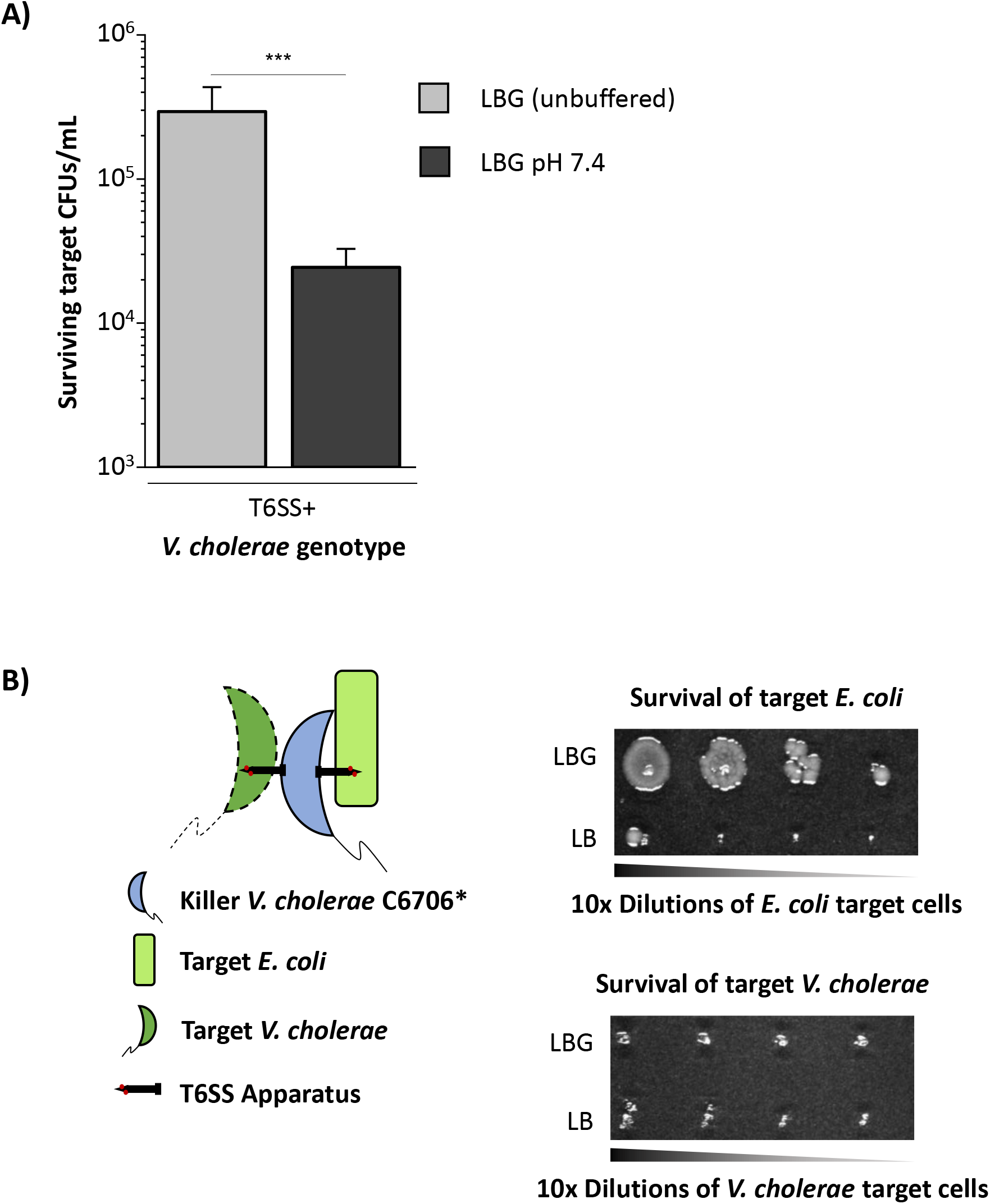
Buffered LBG media at a pH of 7.4 increases killing and glucose does not impair T6SS activity. A) Target *E. coli* MG1655 and killer *V. cholerae* C6706* were co-cultured on either unbuffered LBG medium or buffered LBG medium at a pH of 7.4. A Student’s t-test was performed to determine significance. *** p < 0.001. B) Killer T6SS+ *V. cholerae* C6706* was co-cultured with both *E. coli* MG1655 and susceptible *V. cholerae* target cells.

To further confirm that individual *E. coli* cells survive against *V. cholerae* on LBG, we used confocal microscopy to image the co-culture between sfGFP-labelled *E. coli* MG1655 and unlabeled *V. cholerae* C6706* over time. *E. coli* cells were gradually eliminated when the co-culture was performed on LB medium (Fig. 4, top panels). By contrast, on LBG, many clusters of *E. coli* cells persisted after 180 minutes (Fig. 4, bottom panels). Taken together, these results suggest that individual *E. coli* cells survive T6SS attacks on LBG.

**Figure 4.**
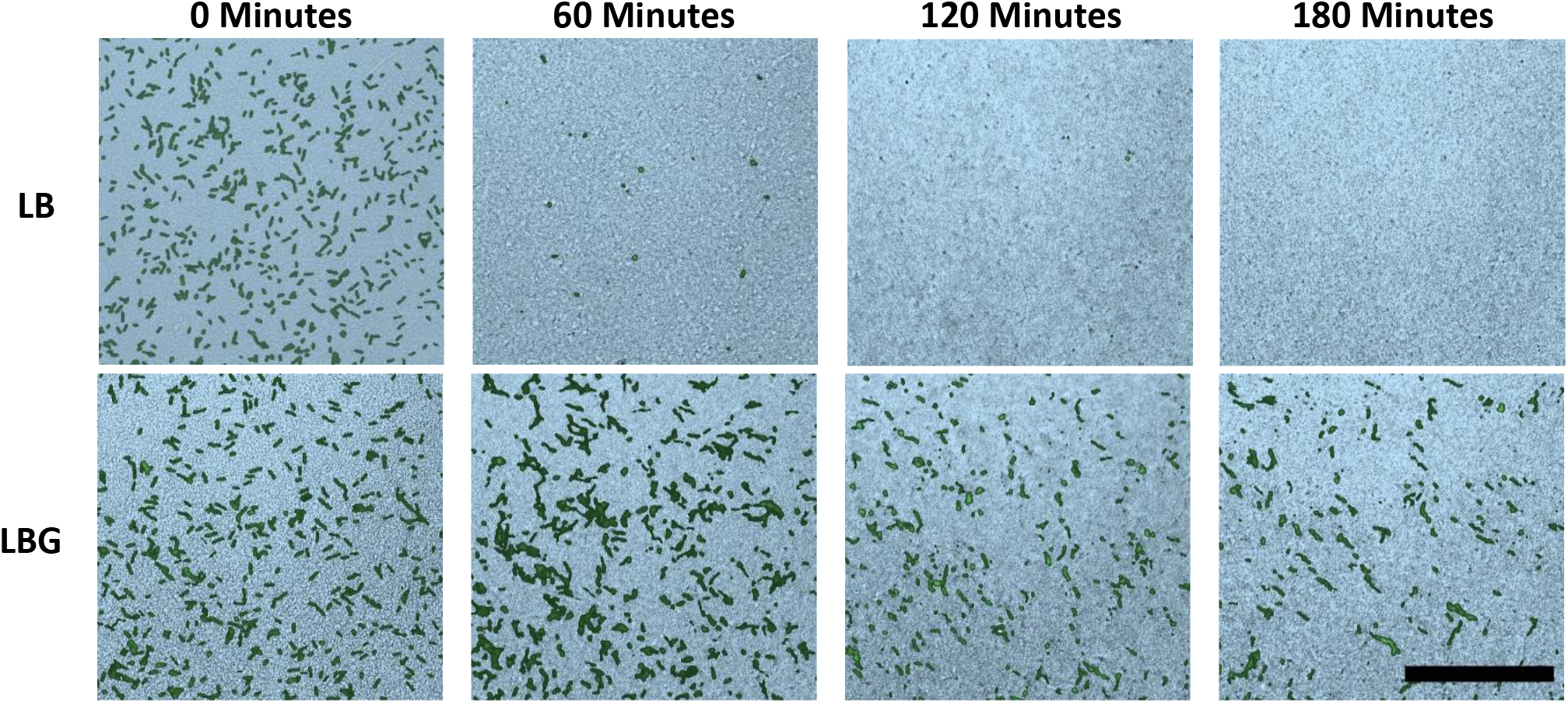
Confocal microscopy confirms that *E. coli* MG1655 cells are resistant to *V. cholerae* T6SS attacks when co-cultured on LBG. Fluorescently labeled green *E. coli* MG1655 cells were densely co-cultured with unlabeled *V. cholerae* C6706* on LB (top panels) or LBG (bottom panels) and imaged for three hours using confocal microscopy. The pseudocolor images depict the brightfield microscopy image (bright blue) of the dense biofilm overlaid with the fluorescence image of sfGFP labeled *E. coli* at different time points. Scale bar is 50 μm.

### A *crp* disruption allows *E. coli* to survive on LB in the absence of glucose

We also investigated whether the *E. coli* resistance is mediated by a physiological response to glucose levels. The cyclic adenosine monophosphate (cAMP) receptor protein (CRP) is a transcriptional regulator that controls expression of hundreds of *E. coli* genes when bound by cAMP during low glucose conditions (42). We disrupted the *crp* gene with a spectinomycin resistance cassette in target *E. coli* MG1655 cells (*E. coli crp*). When co-cultured with *V. cholerae* C6706* T6SS+ on LB lacking glucose, the survival of *E. coli crp* cells was comparable to that of wild type *E. coli* cells when co-cultured with *V. cholerae* on LBG (Fig. 5A). Complementation with a plasmid expressing *crp* restored susceptibility to T6SS attacks (Fig. 5A). To confirm that a *crp* disruption confers protection to *E. coli* on LB, we again used confocal microscopy to observe the co-culture on LB medium. Over the duration of 180 minutes, distinct small domains of sfGFP-labelled *E. coli crp* cells persisted when co-cultured with *V. cholerae* C6706* T6SS+ (Fig. B). Finally, consistent with previous results on LBG, C6706*::Aux 5a bypassed resistance and killed *E. coli crp* on LB in the absence of glucose (Fig. 5C).

**Figure 5.**
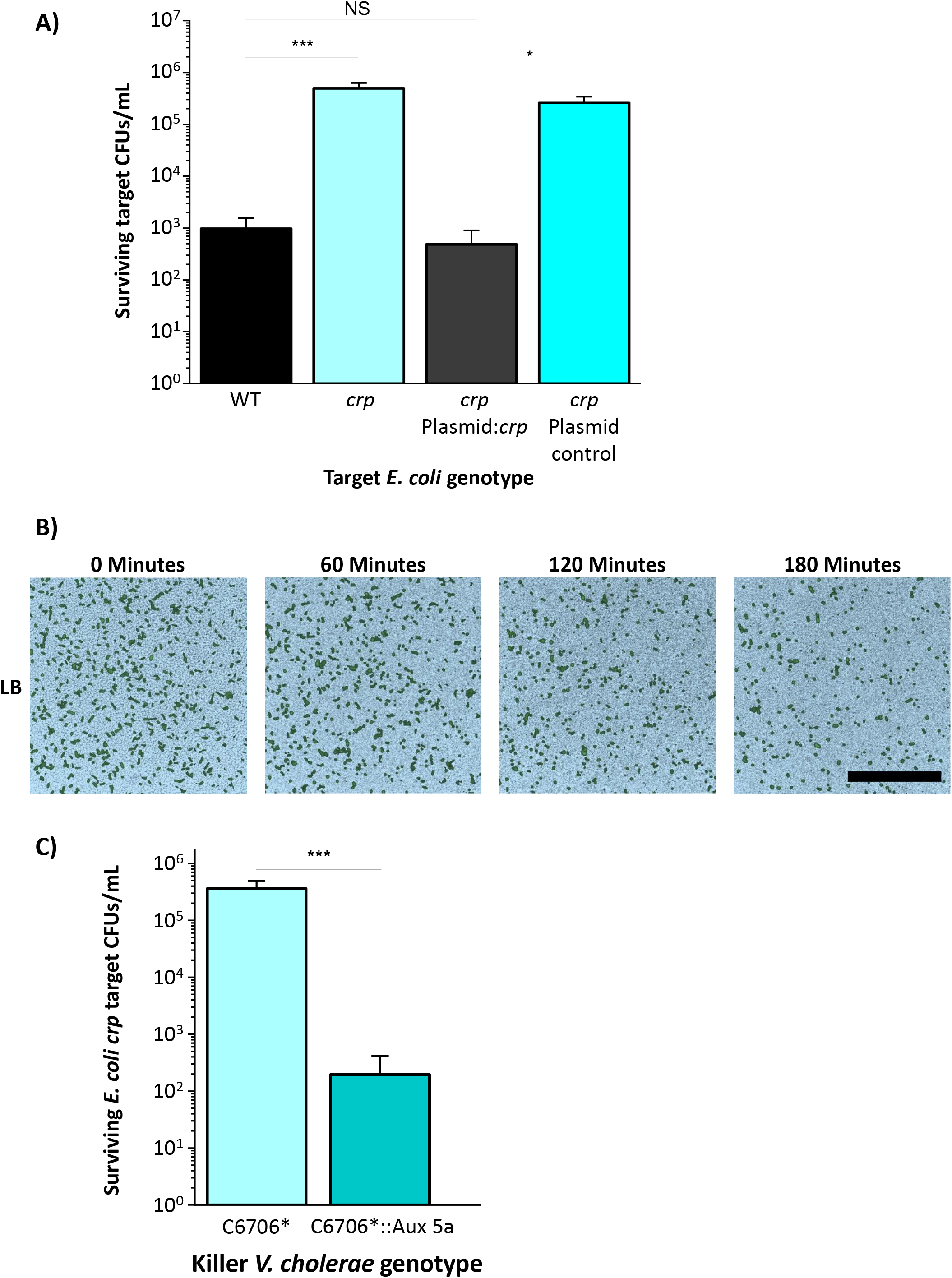
A *crp* disruption confers resistance to *E. coli* MG1655 cells in the absence of glucose. A) *E. coli* MG1655 cells with a *crp* disruption harboring the indicated plasmids were co-cultured with T6SS+ *V. cholerae* C6706* on LB medium. A One-Way ANOVA with a post-hoc Tukey HSD test was used to determine significance. *** p < 0.001, * p < 0.05. NS not significant. B) Fluorescently labeled green *E. coli* MG1655 *crp* cells were densely co-cultured with unlabeled *V. cholerae* C6706* on LB and imaged for three hours using confocal microscopy. Brightfield images (bright blue pseudocolor) were overlaid with fluorescence images of sfGFP labeled *E. coli*. Scale bar is 50 μm. C) *V. cholerae* C6706* or *V. cholerae* C6706*:: Aux 5a were co-cultured with *E. coli* MG1655 *crp* on LB medium. A Student’s t-test was performed to determine significance. *** p < 0.001.

## DISCUSSION

Bacteria live in complex polymicrobial communities with limited resources and fight an evolutionary arms race with other prokaryotes, eukaryotes and viruses (43–45). To succeed in these environments, bacteria have evolved competitive mechanisms including the secretion of antibiotics and contact dependent killing via the T6SS. While those mechanisms represent potent weapons for eliminating competitors, cells have also evolved to encode antibiotic resistance genes and T6SS immunity proteins that neutralize foreign effectors (46–48). Additionally, bacteria face an evolutionary arms race with phages (49). Even though CRISPR-Cas genes protect against viral infections, phages have evolved strategies to bypass bacterial CRISPR-mediated degradation (50–52). Therefore, it is reasonable to hypothesize that an arms race also exists between killer and target bacterial cells leading to the emergence of diverse strategies to defend against the T6SS (18, 53, 54).

In general, cells lacking immunity proteins for cognate effectors are susceptible to T6SS attacks, but studies found that factors other than immunity proteins can also confer protection. McNally et al. have shown that *V. cholerae* cells capable of mutual antagonism spatially segregate into distinct domains that protect target cells at the interior (53). Likewise, large established *E. coli* colonies can persist against T6SS attacks from *V. cholerae* (54). When killer *V. cholerae* cells are co-cultured with target susceptible *V. cholerae*, secreted exopolysaccharides by the target strain also reduce T6SS-mediated killing (15). *P. aeruginosa* cells sense and respond to *V. cholerae’s* T6SS attacks but are not harmed (16). Membrane stress regulators and reactive oxygen species increase resistance of target cells in response to T6SS attacks, while the DsbA disulfide oxidoreductase is necessary for effector activity in target cells (17–20, 55). Colanic acid, an exopolysaccharide secreted by *E. coli* MG1655 when the RcsA protein is overexpressed, protects some strains against phage infection, contact-dependent inhibition and T6SS attacks (18, 55–57). Similar to Hersch et al., we found that *E. coli* MG1655 overexpressing the *rcsA* gene had a mucoid phenotype and displayed a modest increase in survival against *V. cholerae* C6706* T6SS killing on LB (Supp. Fig. 3) (18). Considering that the *rcsA* gene was overexpressed from a multi-copy plasmid and that the survival increase was several orders of magnitude lower than the survival induced by glucose, production of colonic acid is unlikely to be the mechanism by which *E. coli* MG1655 cells resist *V. cholerae* C6706* T6SS attacks on LBG (Supp. Fig. 3, Fig. 1).

Here we demonstrated that human commensal *E. coli* strains respond to physiologically-relevant levels of exogenous glucose and become resistant to T6SS attacks by *V. cholerae* C6706* (37, 56, 57)*. E. coli* growth rates on LB or LBG were not significantly different during a 3-hour monoculture growth assay (Supp. Fig. 1). Therefore, it is unlikely that the *E. coli* resistance against T6SS attacks on LBG can be attributed to the formation of large domains that outnumber *V. cholerae* cells as reported previously (54). This conclusion is also supported by our confocal microscopy results, where we observed that small domains of *E. coli* cells persist even when surrounded by *V. cholerae* cells (Fig. 4).

Previously, glucose has been shown to alter interactions between pandemic *V. cholerae* strains and other bacteria that release acidic compounds in the presence of glucose (24, 25). In zebrafish hosts, *E. coli* cells inhibit growth of *V. cholerae* by secreting acids in the presence of glucose (25). Consistent with previous results where acid-induced growth inhibition of *V. cholerae* occurred only after more than three hours of co-culture with an acid-producing bacterial species, we did not observe a significant decrease in the survival of *V. cholerae* cells when co-cultured with *E. coli* on LBG medium compared to LB (Fig. 1B) (24, 25). In a polyculture assay where *V. cholerae* C6706* cells were incubated with both target *E. coli* MG1655 and susceptible target *V. cholerae* cells, killer *V. cholerae* C6706* cells successfully eliminated target *V. cholerae*, while *E. coli* resisted attacks (Fig. 3B). This result supports a model that the T6SS apparatus is still active in killer *V. cholerae* C6706* T6SS+ cells on LBG. Furthermore, it also shows that the resistance mechanism by which *E. coli* cells survive is not mediated by a product that diffuses over large distances. Taken together, these results suggest that exogenous glucose triggers a response in *E. coli* that allows them to resist T6SS attacks.

Human commensal *E. coli* strains HS, ECOR-2 and Nissle also survived significantly better on LBG media compared to LB, suggesting the glucose-mediated resistance is widespread among diverse *E. coli* strains (Fig. 3A) (31, 33–35, 37). Even though susceptible target *V. cholerae* and *A. veronii* strains displayed an increase in the number of surviving cells on LBG, the resistance was modest compared to *E. coli* strains (Fig. 2A). Future experiments will study the effects glucose has on these and other non-*E. coli* bacterial species, including members of the human gut microbiota.

The glucose-mediated resistance is also dependent on the identity of the killer strain. Clinical strain V52 behaved similarly to C6706* and killed *E. coli* MG1655 to a lesser degree on LBG, although we observed more variation between replicates in the survival of *E. coli* cells when co-cultured with V52 on LBG (Fig. 2C, data not shown). Environmental strains BGT49, BGT64 and BGT69, which encode diverse effectors with activities that are predicted to be different than those encoded by V52 or C6706, efficiently and consistently killed *E. coli* cells on LBG (Fig. 2C) (41). These results suggest that the toxicity of T6SS effectors may be differentially affected by glucose-mediated changes in *E. coli*.

We used a *V. cholerae* C6706* strain engineered to encode the Aux 5a cluster from strain BGT49 and found that the addition of the cluster allowed *V. cholerae* C6706* to bypass the *E. coli* resistance conferred by glucose (Fig. 2C) (41). Furthermore, when *V. cholerae* C6706* cells were co-cultured with *E. coli* MG1655 cells on LBG medium buffered at a pH of 7.4, significantly more killing was observed compared to co-cultures on unbuffered medium (Fig. 3A). The toxicities of some *P. aeruginosa* T6SS effectors are modulated by pH changes while others are unaffected (58). We hypothesize that pH changes induced by the glucose metabolism of *E. coli* cells could also reduce the lethality of T6SS effectors secreted by *V. cholerae* C6706*. We propose that toxic activities of other effectors, like TleV1, are less affected by pH changes and allow killer cells to bypass the glucose-mediated resistance in *E. coli* cells. This could suggest an evolutionary adaptive role for the acquisition of diverse toxins by *V. cholerae* isolates to bypass resistance mechanisms developed in target strains (27, 32, 48). Since *V. cholerae* C6706*:: Aux 5a can kill *E. coli* successfully even when glucose is added to the medium, it is unlikely that effector delivery into *E. coli* cells via the T6SS is prevented on LBG. A recent study found that *E. coli* cells growing in acidic conditions alter their membrane composition by increasing the amount of unsaturated fatty acids (59). This pH-induced membrane change could also affect the toxicity of some *V. cholerae* T6SS effectors. Other factors, including the number of delivered effectors and the ability of killer strains to metabolize glucose might also play roles in bypassing the glucose-induced resistance to T6SS.

The addition of other sugars to LB media resulted in significantly less survival of *E. coli* cells compared to glucose (Fig. 2B). Fructose conferred *E. coli* cells an intermediate resistance against T6SS attacks (Fig. 2B). Glucose and fructose are both recognized and imported into cells using the phosphotransferase system importer (PTS) (60–62). An essential component of the PTS in *E. coli* is EIIA^Glc^ (63). EIIA^Glc^ activates adenylate cyclase when phosphorylated, which leads to an increase in cAMP levels (63). PTS sugars, glucose in particular, cause EllA^Glc^ to become dephosphorylated, resulting in a halt of adenylate cyclase activity and a decrease in cAMP levels (63). cAMP molecules bind and activate CRP, which regulates expression of over 300 genes in *E. coli* (42, 62, 63). Both fructose and glucose PTS transporters have been shown to alter cAMP levels and CRP activity, which could affect expression of genes involved in resistance against T6SS attacks (64, 65).

We found that an *E. coli crp* disruption mutant was resistant to *V. cholerae* C6706* T6SS+ attacks on LB (without glucose induction) in a similar manner to wild type *E. coli* on LBG (Fig. 5A). This indicates that CRP could directly or indirectly repress a gene (or genes) that confers *E. coli* MG1655 protection against T6SS attacks (Fig. 6). When grown on rich media, *E. coli* cells can secrete acetate (66). Yao et al. have shown that tricarboxylic acid (TCA) cycle genes are repressed in *E. coli crp* mutants, leading to a substantial accumulation of acetate (67). We propose that *E. coli* cells with a *crp* disruption could produce sufficiently high levels of acidic compounds on rich LB media, which in turn could lower the pH and neutralize native *V. cholerae* C6706* effectors. Alternatively, CRP could repress stress response genes that protect against T6SS attacks (18, 19, 68).

**Figure 6.**
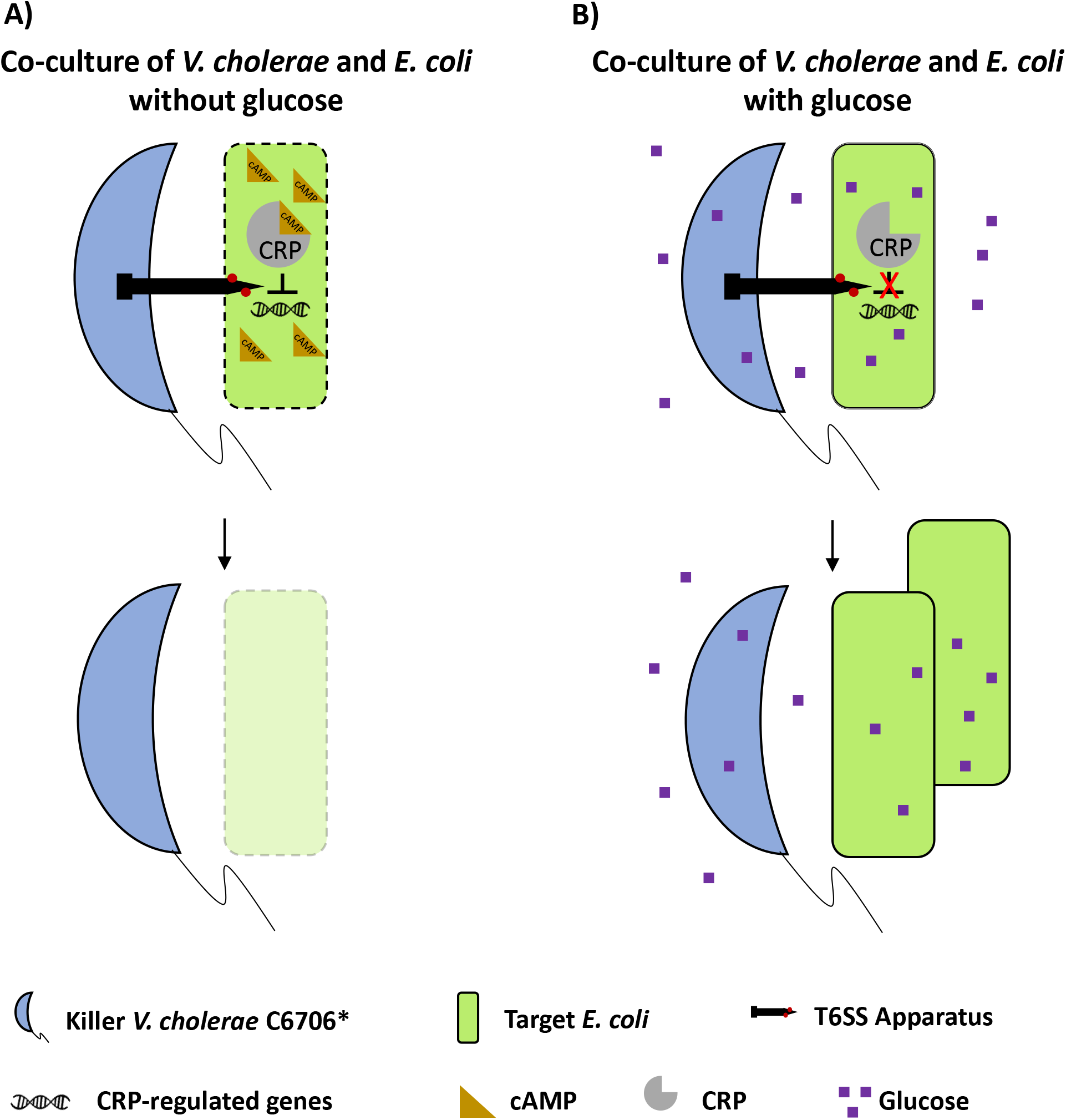
Model depicting the proposed *E. coli* resistance mechanism against the *V. cholerae* C6706* T6SS. A) When *E. coli* cells are grown on rich media in the absence of glucose, CRP is bound by cAMP and represses (directly or indirectly) an unknown gene (or genes). Repression of those genes results in higher susceptibility to T6SS attacks. B) When *E. coli* cells are grown on rich media in the presence of glucose, CRP is not bound by cAMP and cannot repress the genes it controls under low glucose conditions. As a result, *E. coli* cells have increased resistance to T6SS attacks.

To our knowledge, this study highlights the first evidence of an environmentally induced genetic response that increases resistance to T6SS attacks. Zhao et al. have shown that T6SS-mediated killing of *E. coli* cells by *V. cholerae* enhances cholera symptoms in mouse model systems (11). We speculate that metabolites like glucose or other environmental molecules in digestive tracts could also confer resistance to *E. coli* and other commensals to T6SS attacks by other members of the gut microbiota or foreign pathogens. Future studies will be conducted to better understand how *E. coli* and other target cells resist T6SS attacks on glucose, and to determine the role of this mechanism *in vivo*.

## MATERIALS AND METHODS

### V. cholerae and E. coli genetic mutations

All *V. cholerae* mutant strains were made as previously described using the pKAS allelic exchange methods (69). All *E. coli* mutant strains (with the exception of the *E. coli* strain expressing sfGFP) were made as previously described using the Lambda Red system (70, 71). The *E. coli* strain expressing sfGFP was made using an integration event from a pKAS construct. *E. coli* strains Nissle, HS and ECOR-2, as well as *A. veronii* and *V. cholerae* target cells harbored the pSLS3 plasmid to confer chloramphenicol resistance (see Supp. Table 1). Restriction enzymes, polymerases and Gibson mix reagents were used according to the manufacturer’s instructions (Promega and New England Biolabs). Plasmids were verified by PCR and Sanger sequencing (Eurofins).

### Bacterial co-culture assays

Bacterial cultures were grown in liquid LB or LB + glucose 0.4% (LBG) media with shaking at 37°C overnight. Cultures were back-diluted, incubated at 37°C with shaking for 3 hours in the same conditions as the overnight cultures and the absorption for each sample was set to an OD_600_ of 1. Strains harboring plasmids were grown in overnight liquid media with the respective antibiotics required to maintain plasmids and were washed three times with fresh media before co-cultures. Killer and target strains were mixed at a 10:1 (killer:target) ratio and 50 μL of the mixture was spotted on a filter paper with a 0.22 μm pore size. The filter was placed on LB or LBG agar media and incubated at 37°C for 3 hours. Filters were vortexed in 5 mL of sterile LB medium for 30 seconds and 100 μL (or 3μL for spot plating) of serial dilutions were spread on chloramphenicol plates to select for target cells.

Modified co-culture assays were performed as described above for different conditions. Different sugars (fructose, sucrose, lactose, galactose and maltose) were added to LB to a final concentration of 0.4%. Overnight cultures, back-diluted cultures and co-culture experiments were performed on medium containing the respective sugar. Co-culture assays using buffered media were performed on LBG medium adjusted to a pH of 7.4 and buffered with 40 mM MOPS. Experiments were also conducted with buffered LBG medium containing 40 mM HEPES or 100 mM phosphate buffers and similar results were obtained. Overnight cultures, back-diluted cultures and co-culture experiments were performed on pH 7.4 buffered LBG medium. Strains expressing *rcsA* and *crp* were induced with 100 μM IPTG during overnight cultures and co-cultures with killer cells.

### Confocal microscopy

Confocal microscopy experiments were performed as described previously (41). Briefly, overnight cultures were back-diluted 1:100 for 3 hours in the same media as the overnights and the concentration of each sample was set to an OD_600_ of 10. Next, a 1 μL aliquot of a 10:1 killer:target cell mixture was spotted onto an LB or LBG agar pad on a glass slide. Cells were imaged at 96-100% humidity and 37°C using a Nikon A1R confocal microscope. A Perfect Focus System with a 40x objective (Plan Fluor ELWD 40x DIC M N1) was used to stabilize the focus in the plane of the colony growth. Images were processed using ImageJ.

## ACKNOWLEDGEMENTS

We would like to thank Dr. Jacob Thomas for advice and useful discussions, Megan Dillon, Dr. Wai-Leung Ng for providing us the *E. coli* Nissle strain, Dr. Shannon D. Manning for providing us the *E. coli* ECOR-2 strain and Dr. Vanessa Sperandio for providing us the *E. coli* HS strain.

We would also like to thank funding from the Georgia Institute of Technology School of Biological Sciences, NSF MCB 1149925 and the German National Academy of Natural Sciences Leopoldina.

## SUPPLEMENTARY MATERIAL LEGENDS

**Supplementary Figure 1. Glucose does not significantly alter the growth rates of *E. coli* MG1655 or *V. cholerae* C6706* in monoculture.** *E. coli* MG1655 or *V. cholerae* C6706* were grown in monoculture conditions identical to co-culture experiments described in this study. NS not significant. A Student’s t-test was performed to determine significance.

**Supplementary Figure 2. Sugars other than glucose have minimal impact on the recovered number of *E. coli* MG1655 cells.** *V. cholerae* C6706* T6SS-cells were co-cultured with *E. coli* MG1655 cells on LB medium containing 0.4% of the indicated sugars.

**Supplementary Figure 3. Colanic acid confers *E. coli* cell modest protection against *V. cholerae* C6706* T6SS attacks.** *E. coli* cells harboring a plasmid control or a plasmid overexpressing *rcsA* were co-cultured with T6SS+ and T6SS-*V. cholerae* on LB. ** p < 0.01, * p < 0.05.

**Supplementary Table 1. *E. coli* and *V. cholerae* strains and plasmids used in this study.**

